# Cohort Profile: Extended Cohort for E-health, Environment and DNA (EXCEED)

**DOI:** 10.1101/422782

**Authors:** Catherine John, Nicola F Reeve, Robert C Free, Alexander T Williams, Aliki-Eleni Farmaki, Jane Bethea, Linda M Barton, Nick Shrine, Chiara Batini, Richard Packer, Sarah Terry, Beverley Hargadon, Qingning Wang, Carl A Melbourne, Emma L Adams, Catherine E Bee, Kyla Harrington, José Miola, Nigel J Brunskill, Christopher E Brightling, Julian Barwell, Susan E Wallace, Ron Hsu, David J Shepherd, Edward J Hollox, Louise V Wain, Martin D Tobin

**Affiliations:** Department of Health Sciences, University of Leicester, Leicester, UK; NIHR Leicester Biomedical Research Centre, University of Leicester, Leicester, UK; Department of Respiratory Sciences, University of Leicester, Leicester, UK; Department of Population Science and Experimental Medicine, Institute of Cardiovascular Science, University College London, London, UK; Department of Haematology, University Hospitals of Leicester NHS Trust, Leicester, UK; Leicester Law School, University of Leicester, Leicester, UK; Department of Cardiovascular Sciences, University of Leicester, Leicester, UK; Department of Genetics and Genome Biology, University of Leicester, Leicester, UK

## Why was the cohort set up?

EXCEED aims to develop understanding of the genetic, environmental and lifestyle-related causes of health and disease. Cohorts like EXCEED, with broad consent to study multiple phenotypes related to onset and progression of disease and drug response, have a role to play in medicines development, by providing genetic evidence that can identify, support or refute putative drug efficacy or identify possible adverse effects [1]. Furthermore, such cohorts are well suited to the study of multimorbidity.

Multimorbidity describes the presence of multiple diseases or conditions in one patient, though definitions in the literature vary widely [2–4]. It demands a holistic approach to optimise care and avoid iatrogenic complications, such as drug interactions. In the context of increasing specialisation of many healthcare systems and high healthcare utilisation amongst people with multimorbidity, providing such care poses a complex challenge [5–7]. In high-income countries multimorbidity is particularly common amongst more deprived socioeconomic groups and may even be considered as the norm amongst older people [8, 9], whilst an ageing global population and a growing burden of non-communicable diseases in low- and middle-income countries compound its global importance [10]. An expert working group convened by the UK Academy of Medical Sciences recently highlighted the lack of available evidence relating to the burden, determinants, prevention and treatment of multimorbidity, and recommended the prioritisation of research on multimorbidity spanning the translational pathway from understanding of its biological mechanisms to health services research [11].

Studies designed to investigate multimorbidity, rather than considering individual conditions in relative isolation, are therefore vital [6, 7]. Linkage to electronic health records (EHR) has enabled information on a broad range of diseases and risk factors to be studied in EXCEED and places multimorbidity at the study’s heart. The EHR linkage also facilitates longitudinal follow-up over an extended period, enabling, for example, the investigation of lifestyle factors and other exposures on healthy ageing and outcomes in later life.

Combining wide-ranging data from EHR with genome-wide genotyping is also central to EXCEED’s purpose. In recent years, our understanding of which genes are associated with both rare and common diseases has advanced rapidly as available sample sizes for genome-wide association studies (GWAS) have grown rapidly [12]. For example, there are now 279 genetic variants associated with lung function and COPD [13–25]. However, in many cases, our understanding of the mechanisms through which these variants influence disease risk – and which could therefore be therapeutic targets – is relatively limited. An efficient design to inform this understanding is to stratify participants based on available study data on their health status (phenotype) or genetic risk factors (genotype) to recall them for further detailed investigations which would be impracticable across a whole cohort. EXCEED was purposely designed as a resource for recall-by-genotype sub-studies and all participants have consented to be recalled on this basis.

The study is led by the University of Leicester, in partnership with University Hospitals of Leicester NHS Trust and in collaboration with Leicestershire Partnership NHS Trust, local general practices and smoking cessation services.

## Who is in the cohort?

Recruitment to the cohort has taken place since 22nd November 2013, primarily from the general population through general practices in Leicester City, Leicestershire and Rutland. In total, 10 156 participants have been recruited to 4th December 2018. Of these, 445 were recruited through smoking cessation services in Leicester City, Leicestershire and Rutland, 44 through targeted recruitment of those with a recorded diagnosis of COPD in their electronic primary care record, and 117 through additional community-based recruitment focused on Leicester’s South Asian communities (see Figure 1). Whilst a recruitment target of 10 000 has now been reached, community-based recruitment particularly focusing on minority ethnic groups will continue subject to further funding.

**Figure 1.**
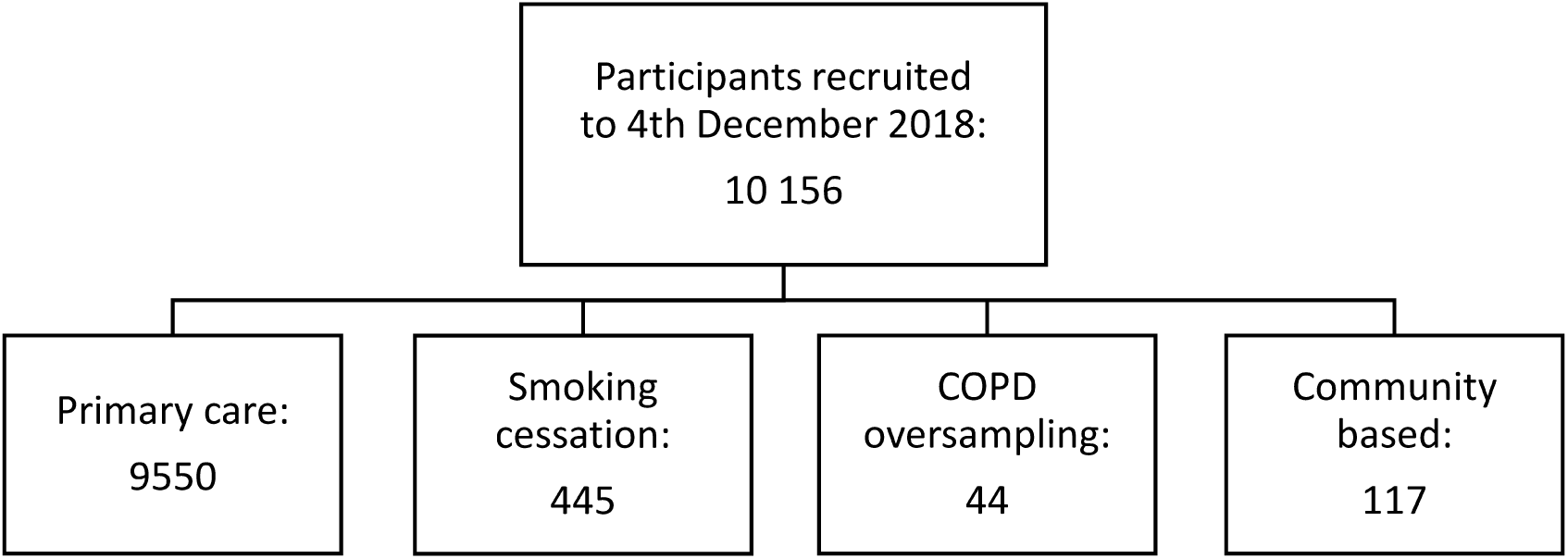
Recruitment methods and numbers

All tables present participants recruited via primary care or smoking cessation whose data was collected and quality control undertaken at 4th December 2018 (9384 participants for questionnaire data, 8930 participants for primary care data). Quality control for the remaining questionnaires and linkage to primary care data is ongoing. Around 400 participants do not have questionnaire data but were recruited as consent and a saliva sample were provided.

In the UK, over 98% of the population is registered with a National Health Service (NHS) general practitioner[26]. For recruitment through primary care, all registered patients aged between 40 and 69 years in participating general practices were eligible for recruitment. Exclusion criteria were minimal: those receiving palliative care, those with learning disabilities or dementia and those whose records indicated they had declined consent for record sharing for research. All eligible patients identified through primary care were sent an initial letter with brief information about the study and a reply slip to indicate their interest.

For participants recruited via smoking cessation services, the lower age limit was reduced to 30 years because of the higher risk of disease amongst smokers. Initial eligibility screening and information provision was either undertaken through electronic client records followed by a letter to the client (as in primary care) or face-to-face by a smoking cessation advisor during a routine appointment. Additionally, patients with a recorded diagnosis of COPD were invited from four local general practices with a higher prevalence of COPD, to boost the numbers available for a sub-study of respiratory disease. For this group, the lower age limit was 30 years, and all other exclusion criteria were identical to the main primary care recruitment.

All those who responded to indicate they were interested in taking part were sent full written information on the study, in addition to a study consent form. Full information regarding participant consent can be found at http://www.leicsrespiratorybru.nihr.ac.uk/our-research/our-research-studies/exceed. All participants consent to follow-up of their electronic healthcare records for up to 25 years, to storage and analysis of their DNA sample, and to being contacted for further studies on the basis of their genetic data (recall-by-genotype) or health status (recall-by-phenotype). They may also consent or decline to be contacted regarding genetic variants which may, in the future, be considered clinically relevant.

Participants proceeded via one of two routes depending on their location and personal preference: a face-to-face appointment with a research professional, or by post. The flow of participants through the main primary care recruitment route is illustrated in Figure 2. Approximately 8% of those who received an initial invitation via primary care completed recruitment. Table 1 gives an overview of the demographic characteristics of the primary care population sampled, compared with the characteristics of those recruited to the study via primary care. Participants in the study were older and more likely to be female than the primary care population from which they were drawn. This reflects well-known patterns of participation in similar cohorts [27, 28]. The local primary care population includes a large proportion of minority ethnic groups, especially Asian and Asian British. These groups are under-represented among study participants, although the proportion of study participants of Asian and Asian British ethnicity (5%) is higher than many UK cohorts, including UK Biobank [28]. This reflects experience of similar recruitment methods in other studies [29].

**Figure 2.**
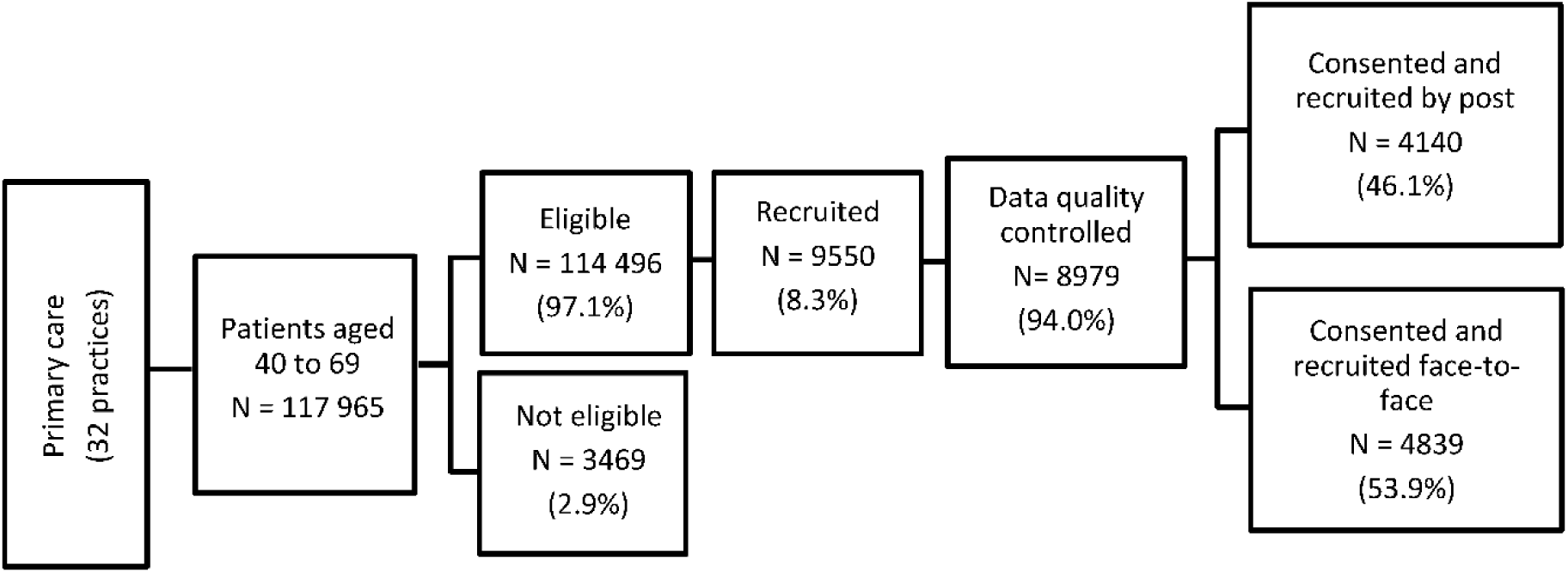
Recruitment via primary care

**Table 1.** Demographic characteristics of the primary care population sampled for the study and those who participated (via the primary care recruitment route only)

|  | Primary care population* |  | Recruited |  | Difference in proportions (95% CI) |
| --- | --- | --- | --- | --- | --- |
| Age | n (N = 117 965) | % | n (N = 8979) | % |  |
| (<45) | 21 057 | 17.9 | 661 | 7.4 | -10.5 (-9.9 to -11.1) |
| (45 – 54) | 44 559 | 37.8 | 2264 | 25.2 | -12.6 (-11.6 to -13.5) |
| (55 – 64) | 36 133 | 30.6 | 3365 | 37.5 | 6.9 (5.8 to 7.9) |
| (≥ 65) | 16 216 | 13.7 | 2689 | 29.9 | 16.2 (15.2 to 17.2) |
| Sex | n (N = 117 965) | % | n (N = 8979) | % |  |
| Male | 59 003 | 50.0 | 3993 | 44.5 | -5.5 (-4.5 to -6.6) |
| Female | 58 962 | 50.0 | 4986 | 55.5 | 5.5 (4.5 to 6.6) |
| Ethnicity | n (N = 81 947) | % | n (N = 8937) | % |  |
| White | 59 576 | 72.7 | 8284 | 92.7 | 20.0 (19.4 to 20.6) |
| Asian/Asian British | 17 670 | 21.6 | 427 | 4.8 | -16.8 (-16.3 to -17.3) |
| Black/African/Caribbean/<br>Black British | 2763 | 3.4 | 12 | 0.1 | -3.3 (-3.1 to -3.4) |
| Mixed | 686 | 0.8 | 93 | 1.0 | 0.2 (0.02 to 0.4) |
| Chinese | 301 | 0.4 | 56 | 0.6 | 0.2 (0.08 to 0.4) |
| Other | 951 | 1.2 | 65 | 0.7 | -0.5 (-0.2 to -0.6) |

\*Primary care population is all patients within the eligible age range in the practices sampled, and includes those who were excluded at the next step (codes for palliative care, dementia, learning disability, or lack of consent to share data for research).

Explanations for the under-representation of minority ethnic groups in medical research more generally include language barriers, inequitable access to healthcare services, cultural sensitivities and a lack of awareness of medical research and its purpose [30, 31]. Community-based recruitment has been introduced to EXCEED to improve representation of these groups.

## How often have they been followed up?

Participants have consented to follow-up through linkage to EHR for up to 25 years. Linkage to electronic primary care records (i.e. records from the participant’s general practice) is undertaken upon completion of recruitment at each practice and has been completed for 8930 participants to the 4th December 2018.

As participants are prospectively followed up we expect losses due to deaths (to date less than 1% of participants), withdrawals (to date less than 0.1% of participants), relatively few losses due to house moves within the UK or changing general practitioner as NHS patients retain the same NHS number and their electronic records move with them, and some losses due to emigration. Analyses of historical healthcare records to track disease development and progression may be subject to selection bias, in particular survivor bias.

## What has been measured?

There are several phases of data collection, summarised in Table 2. Linked primary care data provides historic cohort data. Since the mid-1990s, prospectively recorded consultations enable the retrieval of information not only on symptoms for which participants have visited their general practitioner and diagnoses which have been made, but also on examination findings (including blood pressure readings and spirometry results, for example), laboratory test results, drug prescriptions and secondary care referrals. Major diagnoses documented on paper records prior to the mid-1990s were retrospectively coded at the time of computerisation and so can also be retrieved.

**Table 2.** Summary of data collected at each phase

| Phase | Measurements |
| --- | --- |
| Historic cohort data | Historically coded primary care data (transferred from paper records at the time of practice computerisation, approximately mid-1990s) and since mid-1990s prospectively recorded consultations, with coded:<br>• symptoms;<br>• diagnoses;<br>• measurements, such as blood pressure and spirometry;<br>• laboratory test results;<br>• drug prescriptions;<br>• secondary care referrals. |
| Baseline | All participants:<br>• Questionnaire, including smoking and alcohol use.<br>• DNA saliva sample<br>Postal participants only: self-measured height, weight, waist circumference.<br>Examination by research professional only: height, weight, waist circumference, hip circumference and spirometry |
| Ongoing | Planned quarterly updates to primary care record linkage (detailed above), with consent to follow-up for 25 years, to track health longitudinally<br>Ongoing linkage to:<br>• Admissions, Accident and Emergency attendances and Outpatient appointments via Hospital Episode Statistics<br>• Pathology Data (East Midlands Pathology Service)<br>• Myocardial Ischaemia National Audit (MINAP) |

Baseline data collection for all participants included a self-completion questionnaire which collected detailed information on current and past smoking habits, smoking cessation attempts, e-cigarette and shisha usage, environmental tobacco smoke (second-hand smoke) exposure and alcohol use. For those recruited via a face-to-face appointment, this was undertaken during the appointment. Those participating by post completed the questionnaire online using their own computer, with a paper version available if necessary. Height, weight and waist circumference were either measured by a research professional or self-reported by postal participants. Those recruited face-to-face also had their hip circumference measured and where feasible underwent spirometric measurement of lung function.

Finally, a saliva sample was collected from all participants either at their appointment or returned by post, for extraction of DNA. The samples are stored at the NIHR Biocentre (Milton Keynes, UK), providing industrial-scale laboratory information management and automated robotic systems which have been shown to facilitate efficient error-free sample storage and extraction from freezers in the UK Biobank study [32]. To date, genome-wide genotype data (using the Affymetrix UK Biobank Axiom Array) is available for 5216 participants after quality control, enabling analysis of over 40 million variants after imputation to the Haplotype Reference Consortium (HRC) panel [33].

Planned quarterly updates to linked primary care records will enable longitudinal tracking of health. There is also ongoing linkage to other sources of health data including Admissions, Accident and Emergency attendances and Outpatient appointments via Hospital Episode Statistics; Pathology Data (East Midlands Pathology Service), and the Myocardial Ischaemia National Audit (MINAP).

## What has it found? Key findings and publications

Table 3 shows that, in general, our cohort is slightly healthier than average for common health risk factors and behaviours. This is similar to findings by other cohort studies [28]. For example, the total proportion of participants who were overweight, obese, or morbidly obese (64.2%) was slightly lower than similar age groups in Health Survey for England 2016, where it was above 70% for all ages from 45 upwards [34].

**Table 3.** Prevalence of risk factors and health behaviours

|  | **n** | **%** |
| --- | --- | --- |
| **Deprivation\* (N = 9171)** |  |  |
| 1 (most deprived) | 1204 | 13.1 |
| 2 | 1112 | 12.1 |
| 3 | 1659 | 18.1 |
| 4 | 2507 | 27.3 |
| 5 (least deprived) | 2689 | 29.3 |
| **BMI (N = 9285)** |  |  |
| Underweight (<18.5) | 98 | 1.1 |
| Normal (18.5 – 24.9) | 3221 | 34.7 |
| Overweight (25 – 29.9) | 3576 | 38.5 |
| Obese (30 – 39.9) | 2092 | 22.5 |
| Morbidly obese (≥40) | 298 | 3.2 |
| **Waist circumference (N = 9103)** |  |  |
| Low risk (males<94cm, females<80cm) | 2954 | 32.5 |
| Increased risk (males 94-102cm; females 80-88cm) | 2442 | 26.8 |
| High risk (males>102cm; females>88cm) | 3707 | 40.7 |
| **Smoking status\*<sup>2</sup> (N = 9381)** |  |  |
| Current smoker | 912 | 9.7 |
| Ex-smoker (regular or occasional) | 3678 | 39.2 |
| Never smoker | 4791 | 51.1 |
| **Alcohol intake (units/week) (N = 9335)** |  |  |
| None | 1792 | 19.2 |
| Lower risk (females<14u; males<21u) | 4515 | 48.4 |
| Increasing risk (females 14-35u; males 21-50u) | 2374 | 25.4 |
| Higher risk (females>35u; males>50u) | 657 | 7.0 |

\*Index of multiple deprivation national quintiles by postcode
\*<sup>2</sup>Includes cigarettes, cigars, cigarillos, pipes or shisha

Similarly, the proportion of EXCEED participants who currently smoke is 9.7%, considerably lower than the national average (15.8%) and comparable only to the oldest age group (65 and over) in the national Annual Population Survey, amongst whom smoking prevalence was 8.3%. Smoking prevalence amongst all younger age groups nationally is 15% or above. On the other hand, the proportion of people who report never smoking is also lower than in national population surveys. This may be influenced by question wording and interpretation: while the relevant national survey asked if people had ever “regularly” smoked, the EXCEED questionnaire included occasional use in the definition of ever smokers [35]. Table 4 presents more detailed information on smoking habits amongst current and ex-smokers. The vast majority of both current and ex-smokers reported smoking cigarettes, but cigar/cigarillo and pipe smoking was less common amongst current than ex-smokers. Alcohol intake for our cohort is comparable to that of similar age groups in Health Survey for England 2016 [36].

**Table 4.** Smoking history (self-reported by current or ex-smokers)

|  | Current smokers |  | Ex-smokers |  |
| --- | --- | --- | --- | --- |
|  | n | % | n | % |
| **Type of tobacco used\*<sup>1</sup>** | **(N = 905)** |  | **(N = 3675)** |  |
| Cigarettes*<sup>2</sup> | 857 | 93.6 | 3603 | 97.8 |
| Shisha | 1 | 0.1 | 6 | 0.2 |
| Cigars/cigarillos | 46 | 5.0 | 252 | 6.8 |
| Pipe | 15 | 1.6 | 147 | 4.0 |
| Other | 13 | 1.4 | 2 | 0.1 |
| **Use of electronic cigarettes** | **(N = 909)** |  | **(N = 3675)** |  |
| Ever | 238 | 26.2 | 204 | 5.6 |
| Never | 671 | 73.8 | 3471 | 94.4 |
| **Smoking cessation aids used (ever)\*<sup>3</sup>** | **(N = 377)** |  | **(N = 3636)** |  |
| NRT | 117 | 31.0 | 464 | 12.6 |
| Bupropion | 5 | 1.3 | 40 | 1.1 |
| Varenicline | 54 | 14.3 | 231 | 6.3 |
| Other | 48 | 12.7 | 304 | 8.3 |
| None | 193 | 51.2 | 2655 | 72.2 |
|  | **Mean** | **SD** | **Mean** | **SD** |
| **Pack-years of smoking\*<sup>4</sup>** | **(N = 520)** |  | **(N = 3167)** |  |
|  | 27.2 | 18.9 | 18.4 | 20.6 |
| **Cigarettes per day** | **(N = 562)** |  | **(N = 3197)** |  |
|  | 13.5 | 8.6 | 14.8 | 11.9 |
| **Age at smoking initiation (years)** | **(N = 776)** |  | **(N = 3645)** |  |
|  | 18.4 | 5.8 | 17.1 | 3.8 |

\*<sup>1</sup> People may use more than one type of tobacco, so percentages will not add up to 100.
\*<sup>2</sup> Filtered, unfiltered and handrolled.
\*<sup>3</sup> only for quit attempts lasting at least 6 months. Denominator for percentages is current smokers who have made a quit attempt lasting at least 6 months, or total number of ex-smokers. People may have used more than one aid, so percentages will not add up to 100.
\*<sup>4</sup> only for cigarette smokers

Only 25.2% of participants are in the two most deprived national quintiles and 29.3% are in the least deprived quintile. For Leicester City, 75.9% of the population are in the two most deprived quintiles and only 1.4% are in the least deprived quintile [37]. Though this reflects the whole Leicester population, not just those aged 40-69 and registered with the GP practices that agreed to take part in EXCEED, it indicates that the most deprived communities are under-represented in the cohort.

The Quality and Outcomes Framework (QOF), introduced in 2004, aims to improve the quality of care patients are given by rewarding practices for meeting specified standards of care. Prevalence of 16 chronic conditions prioritised for management in primary care by QOF is presented in Table 5. The figures presented are based on presence of any qualifying diagnostic code [QOF business rules v37, 2017/18] at any time in the patient’s record, with no further exclusions or restrictions. For those conditions where the national QOF prevalence is calculated in a comparable way, prevalence in EXCEED is generally slightly higher and in some cases markedly so. For example, prevalence of hypertension in EXCEED was 28.2% compared to 13.9% nationally. This is likely to be largely due to our older population, since QOF covers all ages.

**Table 5.** Prevalence of chronic conditions

| Condition | n* | % |
| --- | --- | --- |
| Atrial Fibrillation | 243 | 2.7 |
| Asthma | 1138 | 12.7 |
| Cancer | 619 | 6.9 |
| Coronary Heart Disease | 393 | 4.4 |
| Chronic Kidney Disease (3a-5) | 280 | 3.1 |
| Chronic Obstructive Pulmonary Disease | 301 | 3.4 |
| Depression | 2023 | 22.7 |
| Diabetes | 826 | 9.2 |
| Epilepsy | 103 | 1.2 |
| Heart Failure | 107 | 1.2 |
| Hypertension | 2516 | 28.2 |
| Mental Health (psychosis, schizophrenia and bipolar affective disorder) | 71 | 0.8 |
| Osteoporosis | 285 | 3.2 |
| Peripheral Arterial Disease | 51 | 0.6 |
| Rheumatoid Arthritis | 122 | 1.4 |
| Stroke | 108 | 1.2 |

\* Number of participants with one occurrence at any time of a diagnostic code listed in the Quality and Outcomes Framework for that condition. % is out of all participants for whom primary care data was available (8930).

The number of conditions per individual is summarised in Table 6. We found that, overall, 27.2% of our participants had a recorded diagnostic code for more than one QOF condition. This is in line with findings from a large study of almost 100 000 individuals in the Clinical Practice Research Database by Salisbury and colleagues, who used a similar approach to define multimorbidity [5]. They found that 16% of their population had a code for more than one QOF condition, but this rose sharply with age, reaching around 20% amongst 55- to 64-year-olds and over 30% in 65- to 74-year-olds. Two further large UK-based studies have used more comprehensive lists of conditions to define multimorbidity, but limited their focus to active morbidity only, and found prevalence of multimorbidity between 23.2% and 27.2% across all ages, rising substantially with age to 50% or more amongst 65- to 74-year-olds [8, 38].

**Table 6.** Proportion of participants with multiple chronic conditions

| Number of chronic conditions | n | %*<sup>2</sup> |
| --- | --- | --- |
| 1 | 2942 | 32.9 |
| 2 | 1531 | 17.1 |
| 3 | 610 | 6.8 |
| 4 | 218 | 2.4 |
| 5 | 70 | 0.8 |
| 6 or more | 20 | 0.2 |

\*<sup>1</sup>16 chronic conditions prioritised for management in primary care by the Quality and Outcomes Framework (see Table 5)
\*<sup>2</sup> of participants with primary care data

We specifically examined primary care diagnoses of one condition, COPD, for which we had independent diagnostic information from baseline spirometry. Diagnosis of COPD defined by presence of COPD code in primary care data compared with COPD defined by baseline spirometry results indicates that there is underdiagnosis of COPD in EXCEED participants: 84.8% of GOLD stage 1-4 COPD and 71.9% of GOLD stage 2-4 was undiagnosed (Table 7). Existing estimates of the proportion of COPD which is undiagnosed range from around 60% to over 80%, depending on the setting and population studied [39–43]. Using comparable methodology in a similar population to ours, Shahab and colleagues found that 81.2% of those with spirometric COPD had no respiratory diagnosis at all and over 95% had not been diagnosed with COPD.[44] The slightly lower level of underdiagnosis in EXCEED (84.8% of those with spirometric COPD had not received a COPD diagnosis) may be partially attributable to recent improvements in case-finding, and to our use of primary care records rather than self-report to define diagnoses. Reasons posited for such extensive underdiagnosis include a perception amongst clinicians that COPD is solely a disease of elderly smokers, pessimistic views of treatment, lack of availability or underuse of spirometry,[43, 45] and the unreliability of self-reported smoking status in clinical practice.[44]

**Table 7.** Comparison of diagnosis of COPD as defined by COPD codes in primary care data and defined by baseline spirometry

|  |  | COPD defined by baseline spirometry using GOLD criteria |  |  |  |  |  |  |  |
| --- | --- | --- | --- | --- | --- | --- | --- | --- | --- |
|  |  | GOLD 1 – 4 |  |  |  | GOLD 2 - 4 |  |  |  |
|  |  | Yes |  | No |  | Yes |  | No |  |
|  |  | n | % | n | % | n | % | n | % |
| COPD code in primary care | Yes | 86 | 15.2 | 19 | 0.7 | 76 | 28.1 | 29 | 1.0 |
|  | No | 479 | 84.8 | 2643 | 99.3 | 194 | 71.9 | 2928 | 99.0 |

\*For participants with linked primary care data and baseline spirometry measures (n=3,227). All percentages are column percentages.

To demonstrate the utility of EXCEED for enabling cross-sectional or longitudinal studies of quantitative traits, we examined some of the most common measures available in the primary care data, and the numbers of participants with one or more recordings of these measures (Table 8). Table 9 shows the average values of these measures.[46] For example, 98.0% of participants have two or more recordings of blood pressure and over 90% have four or more recordings. Mean systolic blood pressure was 129.9 (sd 13.9) and mean diastolic blood pressure was 78.3 (sd 8.6) (Table 9).

**Table 8.** Numbers of participants with multiple occurrences of a Read code for a quantitative measurement

|  | ≥1 record |  | ≥2 records |  | ≥3 records |  | ≥4 records |  |
| --- | --- | --- | --- | --- | --- | --- | --- | --- |
|  | n | % | n | % | n | % | n | % |
| Blood pressure reading | 8895 | 99.6 | 8748 | 98.0% | 8487 | 95.0% | 8093 | 90.6% |
| Serum creatinine | 8422 | 94.3 | 7492 | 83.9% | 6468 | 72.4% | 5599 | 62.7% |
| Serum sodium | 8417 | 94.3 | 7477 | 83.7% | 6447 | 72.2% | 5582 | 62.5% |
| Serum potassium | 8389 | 93.9 | 7435 | 83.3% | 6400 | 71.7% | 5546 | 62.1% |
| Serum urea level | 8415 | 94.2 | 7460 | 83.5% | 6414 | 71.8% | 5537 | 62.0% |
| eGFR* | 8227 | 92.1 | 7045 | 78.9% | 5883 | 65.9% | 4946 | 55.4% |
| Serum triglyceride levels | 8389 | 93.9 | 6950 | 77.8% | 5633 | 63.1% | 4669 | 52.3% |
| Serum cholesterol level | 8287 | 92.8 | 6852 | 76.7% | 5586 | 62.6% | 4680 | 52.4% |
| Platelet count | 8054 | 90.2 | 6918 | 77.5% | 5827 | 65.3% | 4873 | 54.6% |
| Serum HDL cholesterol level | 8340 | 93.4 | 6770 | 75.8% | 5411 | 60.6% | 4412 | 49.4% |
| Serum LDL cholesterol level | 8050 | 90.1 | 6415 | 71.8% | 5054 | 56.6% | 4091 | 45.8% |
| Serum bilirubin level | 7074 | 79.2 | 5889 | 65.9% | 4835 | 54.1% | 3989 | 44.7% |
| Haemoglobin A1c level | 6782 | 75.9 | 4800 | 53.8% | 3342 | 37.4% | 2383 | 26.7% |
| Total white blood count | 7991 | 89.5 | 6804 | 76.2% | 5674 | 63.5% | 4731 | 53.0% |
| Eosinophil count | 8030 | 89.9 | 6860 | 76.8% | 5728 | 64.1% | 4761 | 53.3% |

\* Glomerular filtration rate calculated by abbreviated Modification of Diet in Renal Disease Study Group calculation [46]

**Table 9.** Summary of values for selected measures

| **Term** | **n\*<sup>1</sup>** | **%** | **Mean** | **sd** |
| --- | --- | --- | --- | --- |
| Blood pressure reading (systolic, mmHg) | 8895 | 99.6 | 129.9 | 14.0 |
| Blood pressure reading (diastolic, mmHg) | 8895 | 99.6 | 78.1 | 8.8 |
| Serum creatinine level (umol/L) | 8422 | 94.3 | 74.1 | 21.3 |
| Serum sodium level (mmol/L) | 8417 | 94.3 | 140.2 | 2.2 |
| Serum potassium level (mmol/L) | 8389 | 93.9 | 4.4 | 0.4 |
| Serum urea level (mmol/L) | 8415 | 94.2 | 5.7 | 1.6 |
| eGFR*<sup>2</sup> (mL/min/1.73m<sup>2</sup>) | 8227 | 92.1 | 82.4 | 10.5 |
| Serum triglyceride levels (mmol/L) | 8389 | 93.9 | 1.5 | 0.8 |
| Serum cholesterol level (mmol/L) | 8287 | 92.8 | 5.1 | 1.1 |
| Platelet count - observation (x10<sup>9</sup>/L) | 8054 | 90.2 | 252.5 | 64.7 |
| Serum HDL cholesterol level (mmol/L) | 8340 | 93.4 | 1.6 | 0.5 |
| Serum LDL cholesterol level (mmol/L) | 8050 | 90.1 | 2.9 | 0.9 |
| Serum bilirubin level (umol/L) | 7074 | 79.2 | 10.5 | 6.8 |
| Haemoglobin A1c level (%) | 6782 | 75.9 | 5.7 | 0.8 |
|  | **n\*<sup>1</sup>** | **%** | **Median** | **IQR** |
| Total white blood count (x10<sup>9</sup>/L) | 7991 | 89.5 | 6.2 | 5.2-7.4 |
| Eosinophil count (x10<sup>9</sup>/L) | 8030 | 89.9 | 0.16 | 0.10 - 0.24 |

\*Where participants have more than one recording of a measure, the most recent value for each participant was used.
\*<sup>1</sup>Number of participants for whom values were available.
\*<sup>2</sup> Glomerular filtration rate calculated by abbreviated Modification of Diet in Renal Disease Study Group calculation [46]

### Recall-by-phenotype study

Recalling by phenotype (see figure 3) facilitates in-depth study of disease mechanisms, with a reduced risk of bias as with nested case-control studies [47]. One such sub-study has recalled EXCEED participants to take part in a study examining the microbiome in COPD cases and in smoking and non-smoking controls.

**Figure 3.**
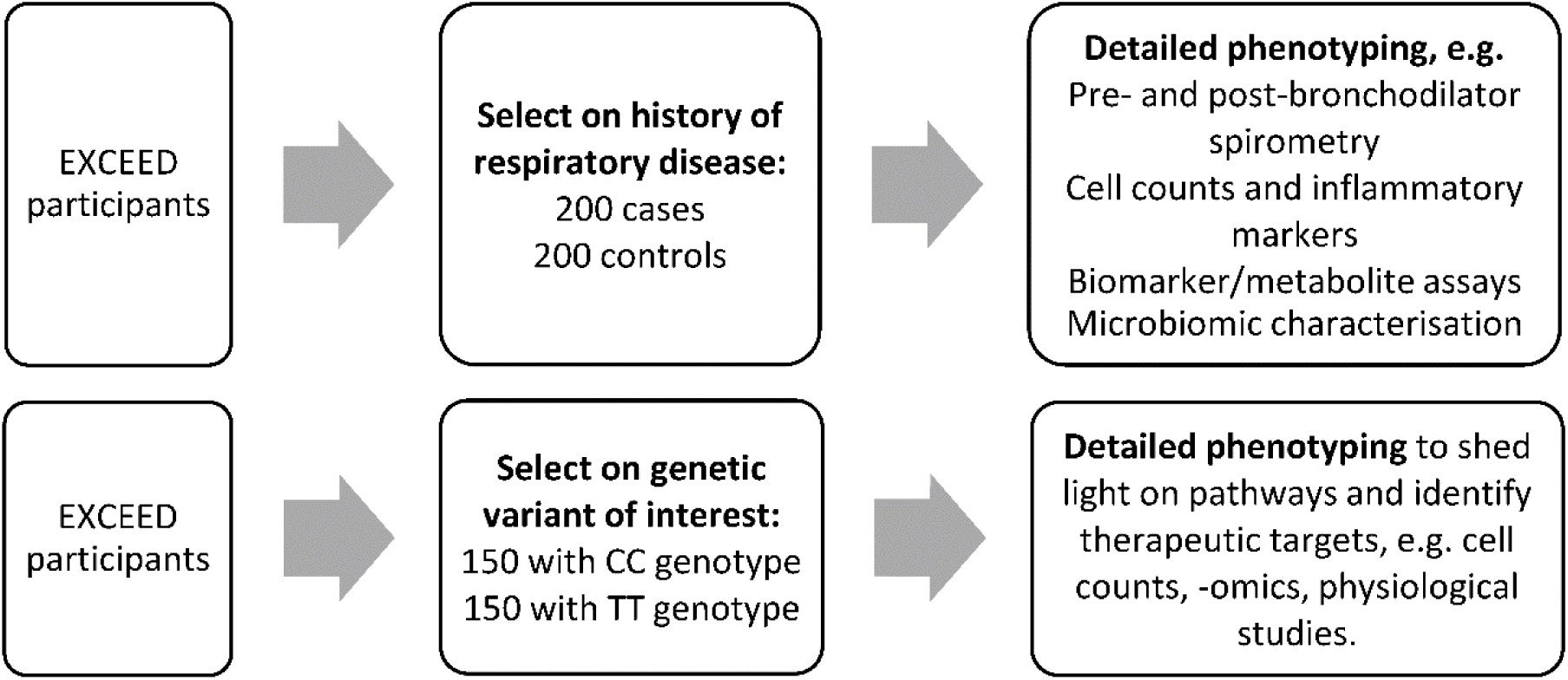
Examples of potential recall-by-phenotype (top) and recall-by-genotype studies (bottom)

### Potential for recall-by-genotype studies

Future recall-by-genotype studies (figure 3) are expected to contribute to a deeper understanding of genetic variants which may be potential therapeutic targets, by bringing back participants for detailed assessments on the basis of the known or suspected mechanism of the relevant gene. Such recall-by-genotype sub-studies may investigate disease susceptibility, disease progression or drug response and whilst they could be interventional in design, most will be observational studies [48]. Observational studies of this kind can provide evidence which is not susceptible to reverse causation and to confounding by lifestyle factors given Mendelian randomisation [49].

Nested designs are also feasible which do not rely on recall of participants but which could be undertaken quickly and inexpensively using stored biological samples and linked electronic data, and such sub-studies could select samples based on either phenotype or genotype. Small-scale intervention-by-genotype studies could, for example, evaluate response to a treatment with a known safety profile in participants with a specific genetic variant.

## What are the main strengths and weaknesses?

Linkage to EHR is a key strength of EXCEED, enabling the study of a wide range of risk factors and diseases, even where data has not been specifically collected at baseline or precedes enrolment as a study participant. UK general practice has had over 20 years of near-universal computerised records [50]. These records have been further enhanced with the introduction of the QOF in 2004, which incentivised GPs to keep comprehensive records of several chronic diseases [51]. Some of these indicators incentivise the recording of quantitative traits relevant to the chronic disease diagnosed, such as blood pressure, lung function, estimated glomerular filtration rate, glycated haemoglobin (HbA1c) and cholesterol measures. That these are expected to be recorded approximately annually means that registered patients often have many repeat measures within linked EHRs, providing an excellent opportunity to study trends in control of conditions such as hypertension or progression of diseases such as COPD. Previous studies have validated some of these primary care measures – for example, routinely recorded spirometry has shown good validity when compared to study specific measures [52]. Other more complex longitudinal outcomes – for example, related to healthy ageing – can also be measured using EHRs.

The use of EHR can have limitations. Misclassification and miscoding of diagnoses may occur, and it is particularly likely that the true prevalence of many diseases will be underestimated (the “clinical iceberg”), as demonstrated by a comparison of COPD diagnoses in primary care data in EXCEED with COPD from spirometry (Table 7). However, the availability of repeat recordings and multiple types of data (including examination findings, pathology results and onwards referrals) over a long period of time can be used to improve and validate the classification of diagnoses and other important exposures and outcomes. Many disease definitions have been validated already – for example, definitions of COPD and asthma in the GOLD-CPRD database – and EXCEED will contribute further to this important area of study [53–59]. In addition to disease status validation, combining records of drug prescriptions, diagnostic and symptom codes can be used define complex phenotypes that have not been possible to study previously.

The cohort recruited adults aged between 30 and 69 years, mostly aged 40 or over, and therefore permits the study of a wide range of questions pertaining to health and disease in adulthood. The absence of younger participants renders it less suitable to study the evolution of disease prior to age 40. This is mitigated by the availability of linked healthcare data from EHR. These records include data prospectively coded by primary care practitioners from the mid-1990s onwards, and will therefore include extensive data from early adulthood for those in middle age when recruited. Earlier life events, for example in childhood and adolescence, are likely to be captured only when they were transferred from paper to electronic healthcare records in the early 1990s, and will therefore include major events such as childhood pneumonia, but not more minor illnesses. The very elderly are also currently absent from the cohort, but data will become available through follow-up of those recruited at the older end of the age spectrum. More generally, the cohort’s age, sex and ethnicity distribution will influence generalisability of research findings to other population groups, and for some research questions validation in other cohorts may be required.

Minority ethnic groups, notably Leicester’s Asian and Asian British population, are currently under-represented in EXCEED. This reflects the recruitment methods utilised to date. We have extended recruitment to increase ethnic minority participant numbers and have adapted our recruitment methods to achieve this, for example, by undertaking recruitment at community events. Minority ethnic groups are also substantially underserved in the availability of samples with genome-wide genotype data worldwide. Whilst the situation has improved in recent years for Asian populations, only 14% of individuals included in genome-wide association studies worldwide up to 2016 were from Asian backgrounds [60]. This situation is replicated in UK-based studies. In UK Biobank, only 2% of participants are from Asian or Asian British ethnic groups, despite this group representing around 7% of the UK population. It is essential that representation of minority ethnic groups increases substantially in genomic studies if these communities are to realise the benefits of genomically–informed advances in precision medicine. EXCEED aims to contribute towards this important goal.

The utility of combining EHR and genetic data for efficient and flexible genetic studies has been highlighted by the eMERGE network of biobanks and Geisinger MyCode [61, 62]. The comprehensive nature and near-universal coverage of NHS health records adds further strength to this study design. In particular the ability to capture virtually all primary and secondary care contacts over decades of the lifespan enables longitudinal studies with a depth of data available in relatively few studies.

Strengths of the study also include consent from all participants to be contacted to participate in recall-by-genotype studies, a type of consent which is not yet widely sought in cohort studies. Recall-by-genotype studies are expected to be highly valuable to identify and validate drug targets and to inform targeting of therapeutics in a precision medicine approach [48]. Maintaining the engagement of cohort participants is important for such studies. Potential strategies include incentivising and/or reducing barriers to further participation, building a study “community” through study branding, newsletters and events, and efforts to trace participants where contact details have lapsed [63, 64]. In EXCEED, in addition to a study newsletter, we are devising approaches with our patient and public involvement (PPI) group, including planned focus groups on dynamic consent approaches.[65]

Some studies incorporating genetic analyses (such as Genomics England) actively seek clinically actionable variants, whilst most cohort studies may not seek to identify these but may discover them as incidental findings. Anticipating this potential, at the time of consent, we asked whether participants would wish to be notified about clinically actionable variants; 99.5% of participants stated that they would wish to be informed in this situation. Clinically actionable variants will be discussed with the regional clinical genetics department of University Hospitals Leicester NHS Trust and then reported back to participants on request for NHS validation. Understanding the reasons for participants’ preferences, how these change over time and how these can best be supported by future policies and procedures will be of key importance for EXCEED and other longitudinal cohort studies.

## Can I get hold of the data? Where can I find out more?

Participants have consented to their pseudonymised data being made available to other approved researchers and we welcome requests for collaboration and data access. Access to the resource requires completion of a proposal form, including a lay summary of the proposed research. Applications to access the resource will be assessed for consistency with the data access policy and with the guidance of the Scientific Committee, which has participant representation. Access to the data will be subject to completion of an appropriate Data/Materials Transfer Agreement and to necessary funding being in place. Requests to collect new data or to utilise biological samples may be subject to additional requirements. Interested researchers are encouraged to contact the study management team via.

### Profile in a nutshell

- EXCEED is a longitudinal population-based cohort which facilitates investigation of genetic, environmental and lifestyle-related determinants of a broad range of diseases and of multiple morbidity through data collected at baseline and via electronic healthcare record linkage.
- Recruitment has taken place in Leicester, Leicestershire and Rutland since 2013 and is ongoing, with 10 156 participants aged 30-69 to date. The population of Leicester is diverse and additional recruitment from the local South Asian community is ongoing.
- Participants have consented to follow-up for up to 25 years through electronic health records (EHR).
- Data available includes baseline demographics, anthropometry, spirometry, lifestyle factors (smoking and alcohol use) and longitudinal health information from primary care records, with additional linkage to other EHR datasets planned. Patients have consented to be contacted for recall-by-genotype and recall-by-phenotype sub-studies, providing an important resource for precision medicine research.
- We welcome requests for collaboration and data access by contacting the study management team via.

## Acknowledgements

The EXCEED study gratefully acknowledges the support of all participants and staff who have contributed to the study. We particularly acknowledge the contribution of the late Dr Corinne Camilleri-Ferrante, Director of Recruitment at the time of the study’s launch.

This paper presents independent research funded partially by the National Institute for Health Research (NIHR) and the Leicester Biomedical Research Centre (BRC). The views expressed are those of the author(s) and not necessarily those of the NHS, the NIHR or the Department of Health.

The study has been supported by the University of Leicester, Leicester City Council, the NIHR Leicester Biomedical Research Centre, the NIHR Clinical Research Network East Midlands, and the Medical Research Council (grant G0902313 to MDT) and the Wellcome Trust (grant 202849 to MDT). LVW holds a GSK/British Lung Foundation Chair in Respiratory Research (grant C17-1).

